# Pre-existing T cell-mediated cross-reactivity to SARS-CoV-2 cannot solely be explained by prior exposure to endemic human coronaviruses

**DOI:** 10.1101/2020.12.08.415703

**Authors:** Cedric C.S. Tan, Christopher J. Owen, Christine Y.L. Tham, Antonio Bertoletti, Lucy van Dorp, Francois Balloux

**Author notes:** Co-last authors (Lucy van Dorp, Francois Balloux).

## Abstract

Several studies have reported the presence of pre-existing humoral or cell-mediated cross-reactivity to SARS-CoV-2 peptides in healthy individuals unexposed to SARS-CoV-2. In particular, the current literature suggests that this pre-existing cross-reactivity could, in part, derive from prior exposure to ‘common cold’ endemic human coronaviruses (HCoVs). In this study, we characterised the sequence homology of SARS-CoV-2-derived T-cell epitopes reported in the literature across the entire diversity of the *Coronaviridae* family. Slightly over half (54.8%) of the tested epitopes did not have noticeable homology to any of the human endemic coronaviruses (HKU1, OC43, NL63 and 229E), suggesting prior exposure to these viruses cannot explain the full cross-reactive profiles observed in healthy unexposed individuals. Further, we find that the proportion of cross-reactive SARS-CoV-2 epitopes with noticeable sequence homology is extremely well predicted by the phylogenetic distance to SARS-CoV-2 (*R*^2^ = 96.6%). None of the coronaviruses sequenced to date showed a statistically significant excess of T-cell epitope homology relative to the proportion of expected random matches given the sequence similarity of their core genome to SARS-CoV-2. Taken together, our results suggest that the repertoire of cross-reactive epitopes reported in healthy adults cannot be primarily explained by prior exposure to any coronavirus known to date, or any related yet-uncharacterised coronavirus.

## Introduction

Severe acute respiratory coronavirus 2 (SARS-CoV-2) is a member of a large family of viruses; the *Coronaviridae*, whose members can infect a wide range of mammals and birds (1). Human coronaviruses were first described in the 1960s (2) with SARS-CoV-2 now the seventh coronavirus known to infect humans; joining the epidemic human coronaviruses SARS-CoV-1 (3) and MERS-CoV (4) and the four species of endemic human coronaviruses (HCoVs). Human endemic coronaviruses are associated with mostly mild upper respiratory infections – ‘common colds’ – and include *Coronaviridae* of the *Alphacoronavirus* genera 229E and NL63 and members of the *Betacoronavirus* genera OC43 and HKU1 (5) to which MERS-CoV, SARS-CoV-1 and SARS-CoV-2 also belong. Both SARS-CoV-1 and SARS-CoV-2 fall into a subgenus of the *Betacoronavirus* named the *Sarbecovirus* (6), with approximately 80% identity at the nucleotide level between SARS-CoV-1 and SARS-CoV-2. All human coronaviruses are thought to be zoonotic in origin, though the exact animal reservoirs remain under debate in some cases (7).

SARS-CoV-2 is estimated to have jumped from a currently unknown animal reservoir into the human population towards the end of 2019 (8) giving rise to the pandemic disease Coronavirus disease 2019 (COVID-19). The symptoms associated with COVID-19 range from fully asymptomatic infections and mild disease through to severe respiratory disease with associated morbidity and mortality. Marked disparities exist in individual risk of severe COVID-19 with gender, ethnicity, metabolic health and age all identified as important determinants (9–11). At a between country level, population age structures and heterogeneous burdens in nursing homes explain some but not all of the variation in infection fatality rates (IFRs) between countries (12). Further important contributors may include climatic variables (e.g. temperature and humidity) and associated seasonal correlates (13–15), the choice of non-pharmaceutical interventions put in place, though with a myriad of other possibly unknown contributing factors.

In light of the wide spectrum of symptoms associated to COVID-19, several studies have probed antibody (16–18) or T-cell responses (19–28) in samples from healthy individuals collected prior to the COVID-19 pandemic to test for the presence of pre-existing cross-reactivity to SARS-CoV-2. Collectively, these findings provide evidence for a degree of T-cell cross-reactivity in unexposed individuals in multiple regions of the world. While the source of this cross-reactivity is still not well-defined, at least some of the cross-reactive T-cell epitopes are suggested to derive from exposure to the four endemic human coronaviruses (19,22), which are circulating in most parts of the world prior to the COVID-19 pandemic, typically in seasonal cycles (29). The relative contribution of each of the four HCoVs to T-cell cross-reactivity patterns observed in unexposed individuals remains unclear. Notably, Peng et al. (25) did not find the presence of cross-reactivity in a cohort of 16 unexposed donors. As such, current evidence suggests that prior exposure to HCoVs may play only a modest role in T-cell cross-reactivity to SARS-CoV-2 in unexposed people.

To date, it also remains unclear whether the detected cross-immunity in unexposed individuals translates into differential COVID-19 pathogenesis. The evidence for a mitigating role of recent HCoV infection on COVID-19 susceptibility and symptom severity upon infection remains conflicting (30,31), and HCoV-reactive T-cells in unexposed individuals have been shown to have only low functional avidity (27). Nonetheless there has been speculation that cross-immunity with the ‘common cold’ endemic HCoVs may, in part, explain variation in the COVID-19 case-fatality rate in different parts of the world (32,33) and that the high incidence of common colds in children and adolescents has contributed to their markedly lower risk of severe disease (18). Additionally, the possible unnoticed circulation in the human population of another animal-associated coronavirus, at least in some regions of the world, cannot at this stage be formally ruled out to have contributed to regional heterogeneities in the spread and associated mortality of COVID-19.

In this study, we employed a bioinformatics approach to probe the possible sources of pre-existing T-cell immunity in samples from healthy individuals predating the COVID-19 pandemic. We analysed sequence conservation over the SARS-CoV-2 proteome across the *Coronaviridae*, which involved the construction of a core gene family-wide phylogeny of all coronavirus representatives that have been sequenced to date. We subsequently assessed the homology to endemic HCoVs and other members of the *Coronaviridae* of 177 CD4^+^ and CD8^+^ epitopes identified in healthy unexposed individuals reported by four independent studies. We find that more than half of the reported epitopes (54.8%) did not have detectable homology to any of the endemic HCoVs. Additionally, none of the sequenced members of the *Coronaviridae* could explain a higher proportion of reported epitopes than expected by chance, given the phylogenetic similarity of their core genome to SARS-CoV-2. Our results suggest that prior exposure to coronaviruses does not primarily explain cross-reactivity patterns to SARS-CoV-2 in unexposed individuals. Instead, patterns of pre-existing T-cell cross-reactivity to SARS-CoV-2 seem in line with lifelong exposure to a diverse and heterogenous array of primarily microbial antigens. We anticipate that our findings will facilitate further characterisations of the potential sources of pre-existing T-cell immunity.

## Results

### Conservation analysis across the family-wide phylogeny of *Coronaviridae*

To reconstruct the genomic diversity of the entire *Coronaviridae* family, we extracted a concatenated alignment of core (shared) genes (ORF1ab, S, M, N) from genome assemblies of 2531 coronaviruses and constructed a Maximum Likelihood phylogeny (**Fig 1a, Table S1**). We then decomposed the SARS-CoV-2 proteome (NC_045512.2) into 15-mer peptide sequences overlapping by 14 amino acids and performed protein BLAST searches to determine the homology to protein sequences translated from each of the 2531 coronavirus assemblies isolated from a range of hosts. The proteome-wide homology of 15-mer peptides across the *Coronaviridae* is represented in **Fig. 1b**. At a 40% sequence identity cut-off, SARS-CoV-2 peptide sequences were highly conserved across the family near the C-terminal end of the ORF1ab polyprotein. Representations of alternative homology thresholds (66% and 80%) provide qualitatively similar patterns (**Fig. S1a** and **Fig. S1b**). This region of homology includes the RNA-dependent RNA polymerase (RdRp) (nsp12) and helicase (nsp13) which are known regions of high conservation across the coronaviruses, with the former frequently used as a taxonomic marker (34).

**Figure 1.**
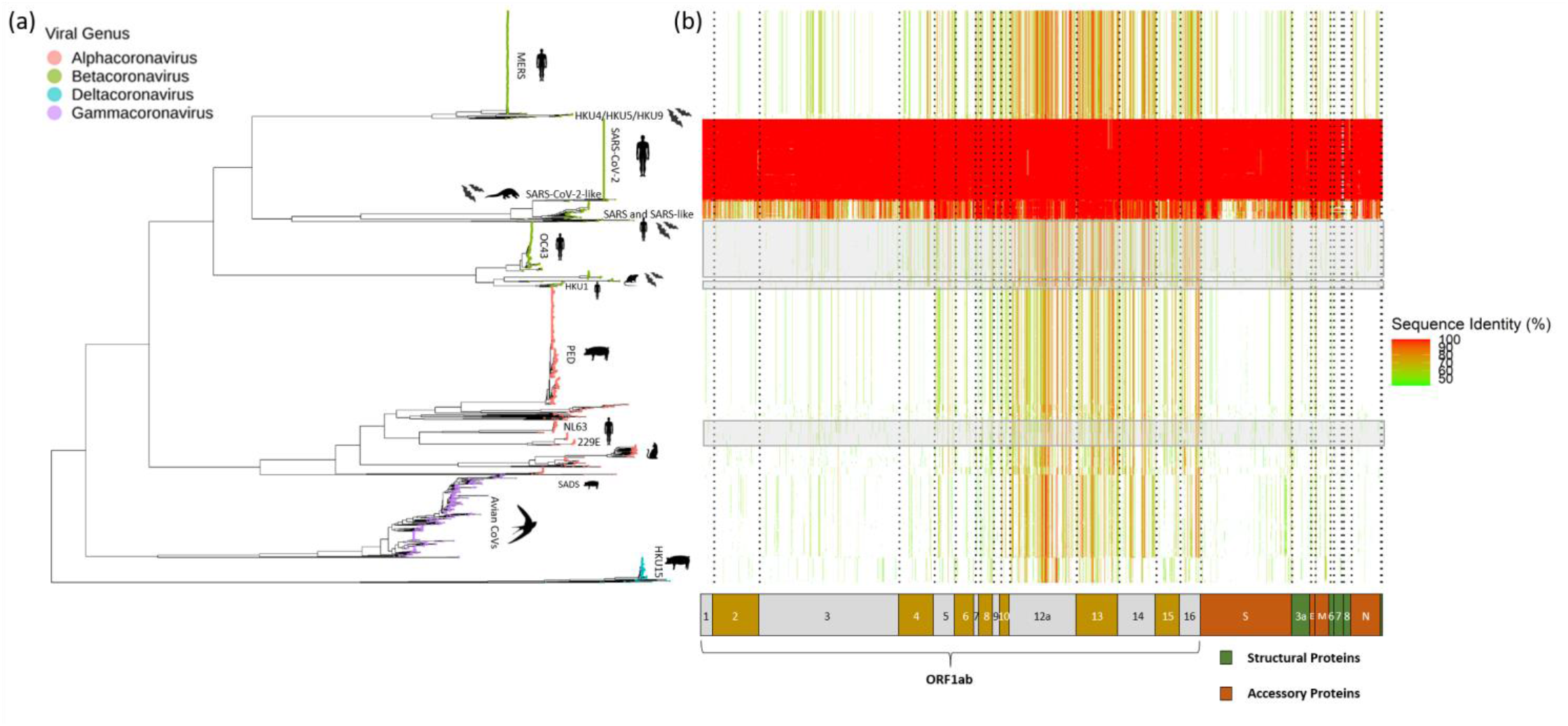
Conservation analysis of SARS-CoV-2-derived 15-mer peptides across the *Coronaviridae*. Maximum likelihood phylogeny of a concatenated alignment of core genes in the *Coronaviridae* annotated by viral genera (tip colour) and highlighting major hosts (**Table S1**). (b) Heatmap visualising the homology of SARS-CoV-2-derived 15-mer peptide sequences across the family. Each row and column correspond to a tip on the phylogeny and a single 15-mer peptide, respectively. The fill of each cell provides the level of homology of a particular SARS-CoV-2-derived 15-mer peptide to the proteome of a single genome record as given by the colour scale at right. Grey boxes highlight the rows of the heatmap corresponding to each of the four endemic human coronaviruses. The homology threshold set to report a protein BLAST hit was 40%.

### Cross-reactivity profiles cannot be completely explained by exposure to endemic HCoVs

We analysed the sequence homology of 177 cross-reactive peptides found to elicit T-cell response in published work on four independent cohorts of healthy unexposed people from Singapore (22), the USA (19) and Germany (23,26) to endemic HCoV protein sequences (**Figure 2**). Notably, we found that 76.3-83.1% of the epitopes could not be explained by homology to any of the four endemic HCoV species individually. In addition, 97 of the 177 epitopes (54.8%) did not have any detectable homology to all the four endemic HCoVs combined (henceforth ‘unexplained’ epitopes). To investigate the potential source of ‘unexplained’ epitopes within the *Coronaviridae* further, we calculated the proportion of these 97 ‘unexplained’ epitopes with detectable homology to each remaining virus in our dataset individually (excluding SARS-CoV-2) (**Figure S2)**. The results suggest that a large proportion of ‘unexplained’ epitopes have detectable homology to at least some of the *Betacoronaviruses* including SARS-CoV-1 and SARS-like coronaviruses within the Sarbecovirus sub-group (**Table S2a**).

**Figure 2.**
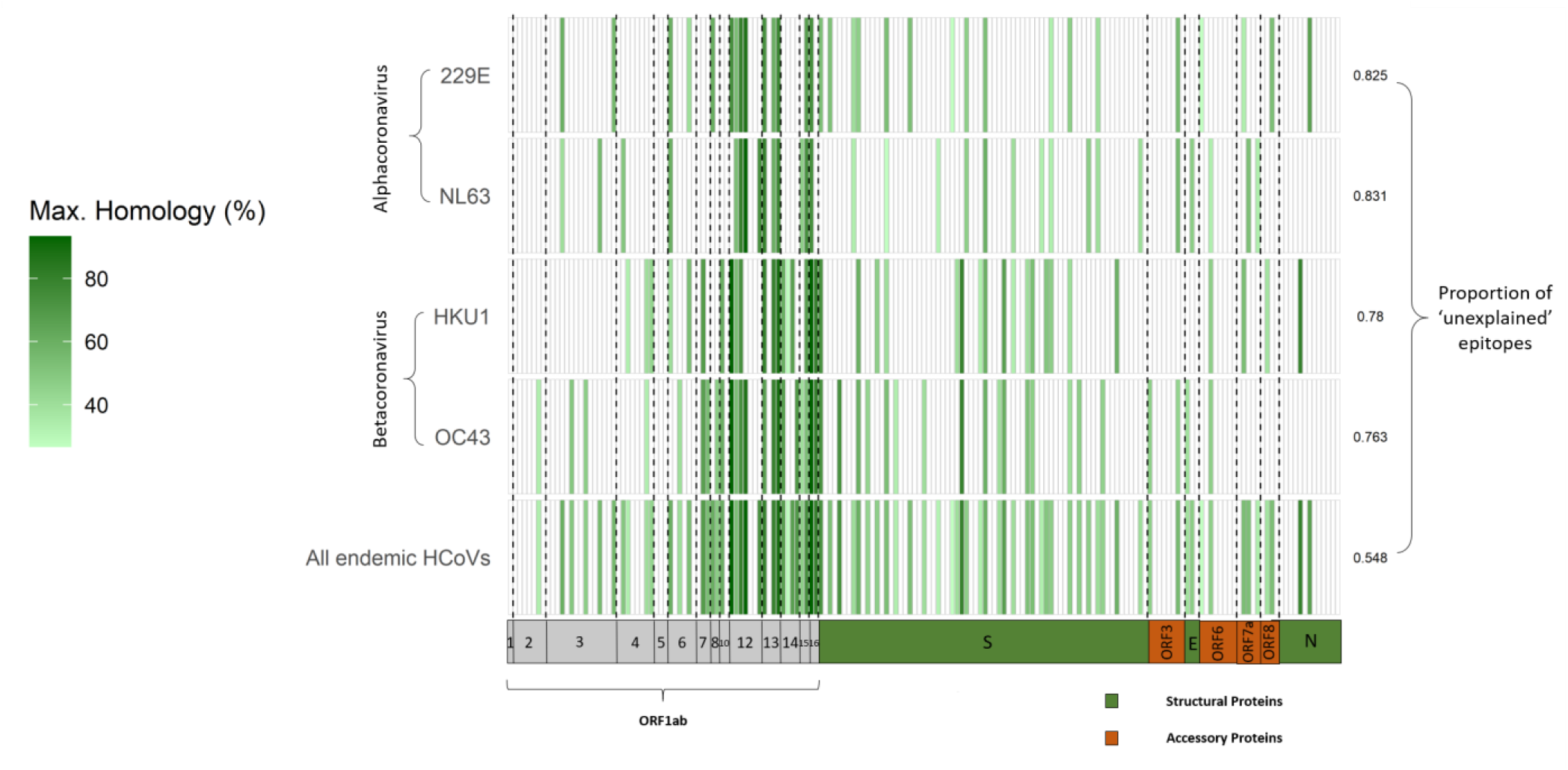
Sequence homology of deconvoluted peptides from published literature to endemic HCoVs. Heatmap visualising the maximum sequence homology of deconvoluted SARS-CoV-2-derived peptides to the each of the four endemic HCoVs (first four rows) and across all HCoVs combined (last row). The proportion of epitopes that cannot be explained by detectable homology to proteins from each species of HCoV is annotated on the right of the heatmap. Each row and column correspond to a single genome record and a single peptide, respectively. The fill of each cell provides the maximum sequence homology of a particular SARS-CoV-2-derived epitope to the proteome of all genome records for each species. This maximum sequence homology was determined by considering only all viruses isolated from a human host and with species names including the terms ‘229E’, ‘NL63’, ‘HKU1’ and ‘OC43’.

Additionally, given the overrepresentation of some species within the dataset, we randomly subset the 2531 viral records to include only one representative of each host and viral species. Using the resultant 155 records, we found that the proportion of published epitopes with detectable homology to coronaviruses is strongly correlated with the natural logarithm of cophenetic distance between each virus relative to SARS-CoV-2 (Pearson’s *r* = -0.983, *p* < 0.0001) (**Figure 3a**). None of the 155 viruses in this filtered dataset had studentised residuals exceeding three, indicating that no coronaviruses within the dataset have homology to a significantly higher number of epitopes than expected by chance (**Figure 3b**).

**Figure 3.**
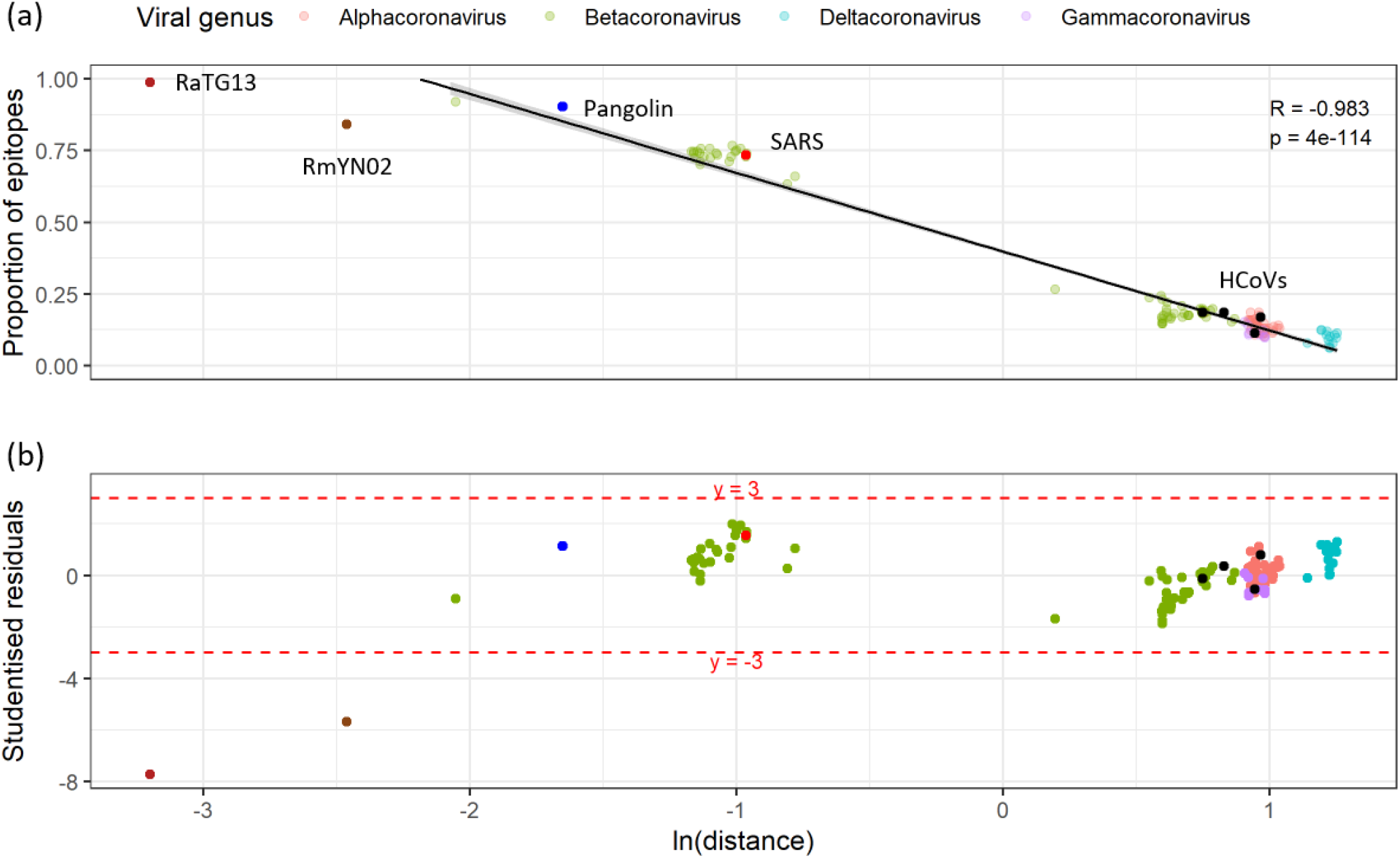
Relationship between the proportion of unexposed epitopes that have detectable sequence homology and the cophenetic distance to SARS-CoV-2 in a representative subset of the *Coronaviridae*. (a) Scatter plot and least squares regression line providing the proportion of epitopes with detectable homology to a coronavirus species (y-axis) and the natural logarithm of cophenetic distance to SARS-CoV-2 (x-axis). The dataset was filtered to only include 155 viruses encompassing all unique host and viral species combinations and are coloured by viral genera, with key members highlighted (**Table S2b**). Pearson’s correlation coefficient and its associated *p*-value of the two variables were calculated using the *cor*.*test* function in *R*. (b) Scatter plot of studentised residuals calculated using the function *studres* from the *MASS* package (35) in *R*.

### Possible sources for T-cell cross-reactivity beyond coronaviruses

To identify possible sources for the T-cell cross-reactivity observed in people unexposed to SARS-CoV-2, we also performed a protein BLAST search for all 177 experimentally validated epitopes against the NCBI non-redundant protein database (excluding the taxon *Coronaviridae*), storing the first 1000 hits in each case. A fraction of the epitopes (10/177) share partial homology with proteins from a very diverse range of taxa, including viruses, bacteria and unicellular eukaryotes (**Table S3**). However, the lowest Expect (E) value of the protein BLAST hits, which represents the number of similar hits expected by chance given the size of the database used and the length of the query (36), is 7.5. This suggests that all the hits shown in **Table S3** could be explained by chance alone. Together with the wide diversity of taxa identified, the results suggest that there is no single candidate for the source(s) of the T-cell cross-reactive repertoire beyond the *Coronaviridae*.

## Discussion

SARS-CoV-2 cross-reactive T-cells in healthy unexposed individuals have been identified as potentially important contributors to the immunological response to COVID-19. Prior exposure to globally circulating endemic coronaviruses present some of the strongest candidates for eliciting such cross-immunity. Though, the relative contribution of these coronaviruses to the reactive T-cell epitopes identified in multiple cohorts of healthy individuals have been only partially explored. We characterised the amino acid homology of SARS-CoV-2-derived T-cell epitopes reported in COVID-19 unexposed individuals from Singapore (22), the USA (19) and Germany (23,26) against the entire proteome of the *Coronaviridae* family, including all major mammalian and avian lineages.

Following a comprehensive screen, we found that 54.8% of reported T-cell epitopes did not have any detectable homology to the four human endemic coronavirus species (HKU1, OC43, NL63 and 229E) (**Figure 2**), despite HCoV infections circulating widely in global human populations (5). We note that the highest conservation to confirmed T-cell epitopes tended to be within members of the *Sarbecovirus* sub-group, which includes SARS-CoV-1, SARS-CoV-2, and a few related species that have been isolated mostly from bats and pangolins but are not known to have been in widespread circulation in humans. However, this homology can be well explained by the phylogenetic affinity of these viral species to SARS-CoV-2 (**Figure 3**). In addition, we note that the region of high sequence homology across all coronaviruses (nsp12-nsp16) (**Figure 1**) is not a primary immune target in COVID-19 convalescent patients (CD8^+^ T-cells). Furthermore, SARS-CoV-2 infection leads to a heterogenous pattern of cell-mediated immune responses over the entire SARS-CoV-2 genome, largely falling outside of the spike protein, not enriched in the terminal end of ORF1ab largely conserved among the coronaviruses, and does not consistently lead to cross-reactivity with endemic HCoVs (37).

Our work adds to a growing suite of evidence that prior HCoV infections are not the sole, and possibly not even the main, candidates responsible for cross-reactive T-cell epitopes in SARS-CoV-2 unexposed individuals. We argue that previous studies that presented empirical evidence of T-cell cross-reactivity with HCoV-derived peptides did not take into account the genetic relatedness of endemic HCoVs to SARS-CoV-2, placing an over-emphasis on these viruses as the source of pre-existing T-cell immunity. This opens the question as to what other antigens may have primed the intrinsic cross-reactivity identified (38) in pre-pandemic samples. A sizeable fraction of cross-reactive T-cell epitopes remains unexplained by prior exposure to any known coronavirus in circulation. It feels fairly implausible that the ‘unexplained’ cross-reactive epitopes are due to prior exposure to a yet undescribed coronavirus. Indeed, such a hypothetical yet-to-be described coronavirus would have needed to be in circulation globally until very recently and then vanished, which seems highly unlikely. Additionally, since we incorporated the whole known genetic diversity of coronaviruses in our analyses, which has been extensively sampled, such an unknown pathogen would have to be phylogenetically unrelated to any coronavirus characterised to date. As such, an unknown coronavirus would be an unlikely candidate for as a source of this ‘unexplained’ T-cell cross-reactivity. Possible alternative agents for the unexplained cross-reactive epitopes may include widespread microbes, or widely administrated vaccines. The tuberculosis bacille Calmette-Guerin (BCG) vaccines have been suggested as candidates providing some cross-immunity against SARS-CoV-2 (39,40). However, our screen of all 177 published T-cell epitopes found no homology to any *Mycobacterium* species (**Table S3**). As such, BCG vaccination represents a most unlikely contributor to the T-cell cross-reactivity observed. Instead we identify a diverse spread of putative antigens with low detectable homology. The presence of such a broad pre-existing repertoire of CD4^+^ reactive T-cells in healthy adults has previously been observed in the context of cross-reactivity to HIV and influenza infection, and interpreted as the result of prior exposure to environmental antigens (41) or proteins in the human microbiome (38). It has also been postulated that the cross-reactive profile may take on an increasing role with age and immunological experience (42) which may result in high levels of inter-individual variation based on infection history and HLA type.

Admittedly, sequence homology is an indirect proxy for probing the source of T-cell cross-reactivity. Yin and Mariuzza (43) reviewed five putative mechanisms of T-cell cross-reactivity, all of which highlight the complex and diverse molecular interactions of peptide, major histocompatibility complex (MHC) and T-cell receptors. In particular, molecular mimicry would suggest that conservation of structure can compensate for lower sequence homology (44–46). At the same time, higher sequence homology improves the likelihood that structural or chemical characteristics are conserved. Deconvolving the relationship between sequence homology and cross-reactivity is evidently non-trivial and remains a limitation of our work. Indeed, we do not rule out the possibility that peptides of lower homology from members of the *Coronaviridae* can result in cross-reactivity. However, we note that the sequence homology analysis of HCoVs and SARS-CoV-2 epitopes by Mateus et al. (19) suggests a positive association of sequence homology and the frequency of cross-reactivity, providing an empirical basis for our approach.

Our results highlight the importance of considering the wider phylogenetic context of circulating antigens contributing to immunological memory to novel pathogens. The widespread and repeated exposure of global human populations to circulating endemic HCoVs is expected to have left an immunological legacy which might modulate COVID-19 pathogenesis. However, our results suggest that the extensive observed T-cell cross-reactivity is unlikely to have been caused by prior exposure to any known coronavirus in global circulation. It is nonetheless clear that the potential cross-reactive repertoire is widespread and present in cohorts of healthy people from multiple countries around the globe (19–28), even if perhaps at low avidity (27). It remains to be established to what extent such cross-reactivity translates into immunity to SARS-CoV-2, both in terms of susceptibility to infection and symptom severity upon infection.

## Methods

### Data acquisition

3300 publicly available complete *Coronaviridae* assemblies were downloaded from NCBI Virus using the *taxid*: 1118 together with accompanying metadata on 08/04/2020. Additionally, we downloaded 12 bat and pangolin Coronavirus sequences from GISAID (47) (acknowledgements in **Table S4**). Sequence duplicates were identified and removed from the combined dataset using *seqkit rmdup* (48) together with those with >10% of sites set to N. Accessions were later retained in the dataset only for those with a reported host of isolation. This resulted in a final dataset of 2533 assemblies with complete metadata with the latter manually cleaned to ensure consistent reporting of host and viral species.

### Maximum Likelihood phylogeny of Coronaviridae

To reconstruct the genomic diversity of the entire *Coronaviridae* family, we extracted the shared core genes from the representative genome assemblies across all genera. First, open reading frames (ORFs) were identified using the genome annotation tool *Prokka* v1.14.6 (49). Next, the *Roary* pipeline v3.11.12 (50) was used to cluster all *Coronaviridae* ORFs at a minimum amino-acid homology threshold of 30%. Sequences for the four genes ORF1ab, S, M and N were each found to cluster in a minimum of 2531 assemblies, which were then extracted, concatenated and aligned using *MAFFT* v7.453 (51). The resulting alignment was trimmed of gaps found in 20% or more isolates and used to build a Maximum Likelihood phylogeny using *RAxML* v8.2.12 (52) with 1000 bootstraps for node support. We provide the curated metadata of the final 2531 viral records used in our analysis in **Table S1**.

As it was not possible to include an outgroup in the *Coronaviridae* concatenated-core alignment, an alignment-free analysis was used to identify the most basal genus with which to root the family Maximum Likelihood phylogeny. All *RefSeq* genome assemblies belonging to the virus order *Nidovirales* were downloaded, which contained 103 sequences accrsoss the sub-orders *Arnidovirineae, Cornidovirineae, Mesnidovirineae, Nanidovirineae, Ronidovirineae* and *Tornidovirineae*. Each assembly contained a ORF1ab CDS annotated ORF, the only gene shared by all members of the *Nidovirales* (53), which were decomposed into 11-mer sequences using *MASH* v2.1.1 (54). Based on pairwise Jaccard Distances of matched 11-mers between all ORF1ab sequences, a Neighbour-Joining tree was constructed to assess the genetic relationship between members of the *Nidovirales*. The genus *Deltacoronavirus* was identified to be the most basal clade of the *Coronaviridae* in the wider context of the taxonomic order and was therefore used to force-root the family Maximum Likelihood phylogeny.

### Sequence conservation analysis

We decomposed the SARS-CoV-2 proteome (sequences retrieved from *RefSeq*; NC_045512.2) into 9394 15-mer peptides overlapping by 14 amino acids using a custom *R* script (https://github.com/cednotsed/tcell_cross_reactivity_covid/blob/main/utils/make_fasta_out_of_proteins. R). In addition, we retrieved the sequences of 177 epitopes found to elicit a response in at least one individual from Singapore (22), the USA (19) and Germany (23,26) from published supplementary tables. The breakdown of the number of epitopes for each T-cell response type is shown in **Table S5b**. Translated protein sequences of all ORFs from each of the 2531 assemblies were retrieved from *Prokka* (49) and used to construct a protein BLAST database. Separately, a protein BLAST database was also constructed from the protein annotations associated with the 2531 assemblies, which were downloaded using *NCBI Batch Entrez* (https://www.ncbi.nlm.nih.gov/sites/batchentrez). Subsequently, we used *blastp* from *BLAST+* v2.11.0 (55) to determine the sequence similarity of the 15-mer peptides from the SARS-CoV-2 proteome and the 177 published epitopes using the two databases and. The resultant protein BLAST outputs were merged by retaining only the hit with the maximum percentage identity for each assembly and query combination. To maximise the number alignments obtained we set *-num_alignments* and *-evalue* parameters to 10^9^ and 2 x 10^9^, respectively. In addition, to optimise the protein BLAST search for short sequences, *-task* was set to *blastp-short*. Lastly, only alignments involving the full length of the query sequence were considered by setting *-qcov_hsp_perc* as 99. This threshold was employed because the query sequences are short and so sequence identity would only be a meaningful measure of homology in alignments given the whole sequence.

### Proportion of published epitopes and cophenetic distance

Using the merged output of the protein BLAST search querying the 177 published epitopes, we analysed the proportion of epitopes that had detectable homology to each virus in a representative filtered dataset of all combinations of unique host and virus species (*n* = 155). The cophenetic distance of each virus relative to SARS-CoV-2 was calculated using *cophenetic*.*phylo* from the *ape* package v5.3 (56) in *R* from the Maximum Likelihood *tree* file. A least squares regression of the proportion of epitopes with detectable homology on the natural logarithm of cophenetic distance was performed using the *lm* function in *R*. Pearson’s correlation of the two variables was calculated using the *cor*.*test* function in R. The studentised residuals were calculated using the *studres* function as part of the *MASS* package v7.3-53 (35).

### Non-*Coronaviridae* protein BLAST

To determine if any proteome outside of the *Coronaviridae* had detectable homology to any of the 177 epitopes reported in the literature, we performed a protein BLAST using the online *blastp suite* (https://tinyurl.com/y22o4t9z) against the non-redundant protein sequence database (accessed 7/12/2020), while excluding sequences associated with the *Coronaviridae* (taxid: 11118). Protein BLAST searches were conducted in eight batches of 20 and a ninth batch of 17 epitopes with the number of alignments performed set to 1000 per batch. After merging the outputs of the eight batches, we filtered the resultant table to exclude missing organism names, hits with descriptions containing the terms ‘synthetic’, ‘SARS’, ‘coronavirus’, or ‘cov’, or organism names labelled as ‘uncultured bacterium’. Additionally, we excluded hits to the accession 6ZGH_A, which contains a region of the SARS-CoV-2 spike protein sequence.

## Data and code availability

All source code used for the analyses can be found on GitHub (https://github.com/cednotsed/tcell_cross_reactivity_covid.git). Genomic data for the *Coronaviridae* were obtained from publicly available accessions on NCBI Virus. The 12 further bat and pangolin associated coronaviruses were also included from the GISAID repository, with full acknowledgements provided in **Table S4**. The list of epitopes used and the frequency table of CD4^+^ and CD8^+^ T-cell epitopes stratified by study cohort can be found in **Table S5a** and **Table S5b** respectively.

## Acknowledgements and Funding

L.v.D and F.B. acknowledge financial support from the Newton Fund UK-China NSFC initiative (grant MR/P007597/1) and the BBSRC (equipment grant BB/R01356X/1). L.v.D. is supported by a UCL Excellence Fellowship. C.O. is funded by a NERC-DTP studentship. Finally, we acknowledge the large number of research groups openly sharing SARS-CoV-2 genomic and immunological data with the research community.

## Supplementary Material

**Figure S1.**
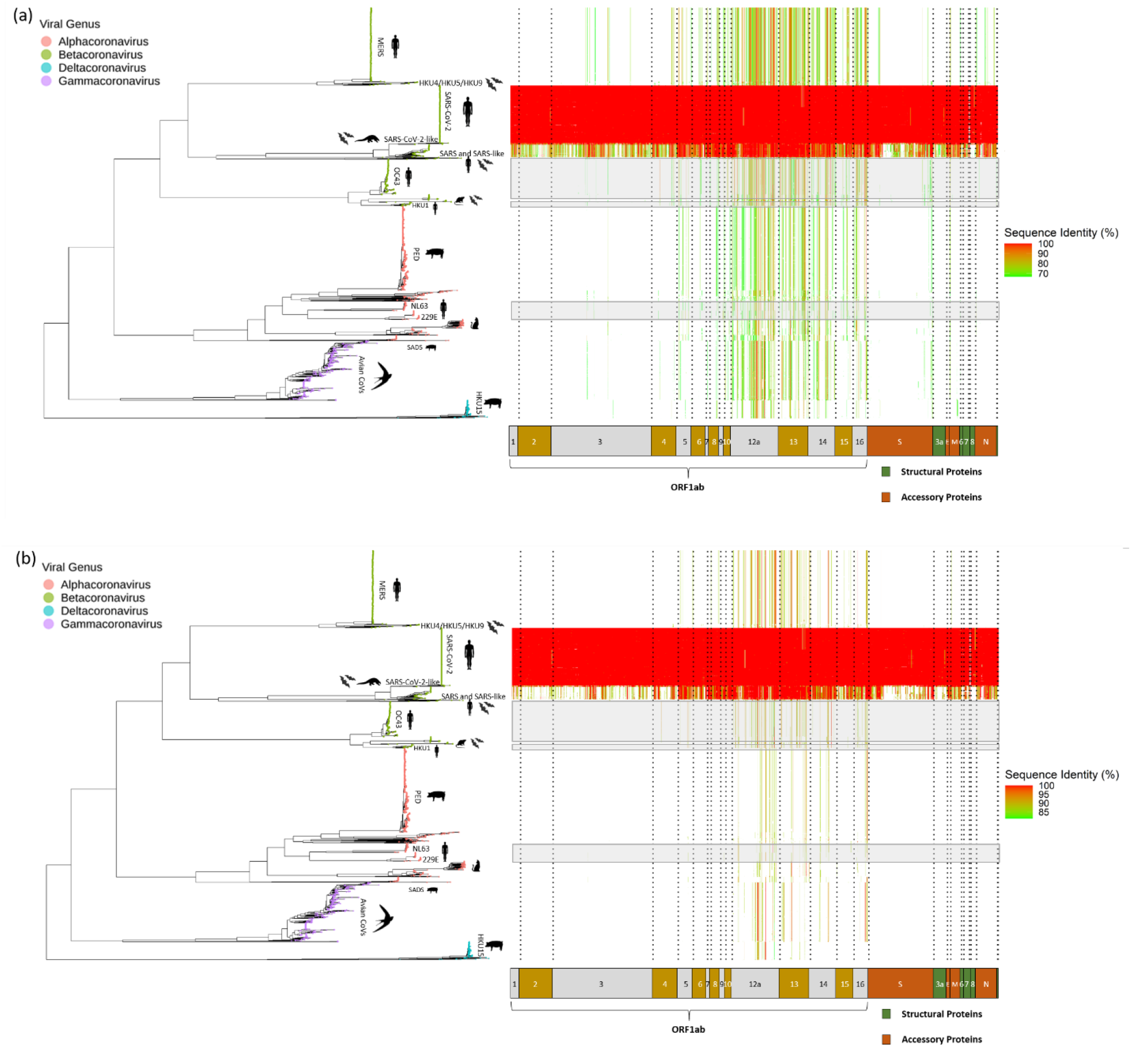
Conservation analysis of SARS-CoV-2-derived 15-mer peptides across the *Coronaviridae*. Maximum likelihood phylogeny and heatmap visualising the homology of SARS-CoV-2-derived 15-mer peptide sequences across the family, similar to that shown in **Figure 1** but using (a) 66% and (b) 80% as the protein BLAST homology threshold.

**Figure S2.**
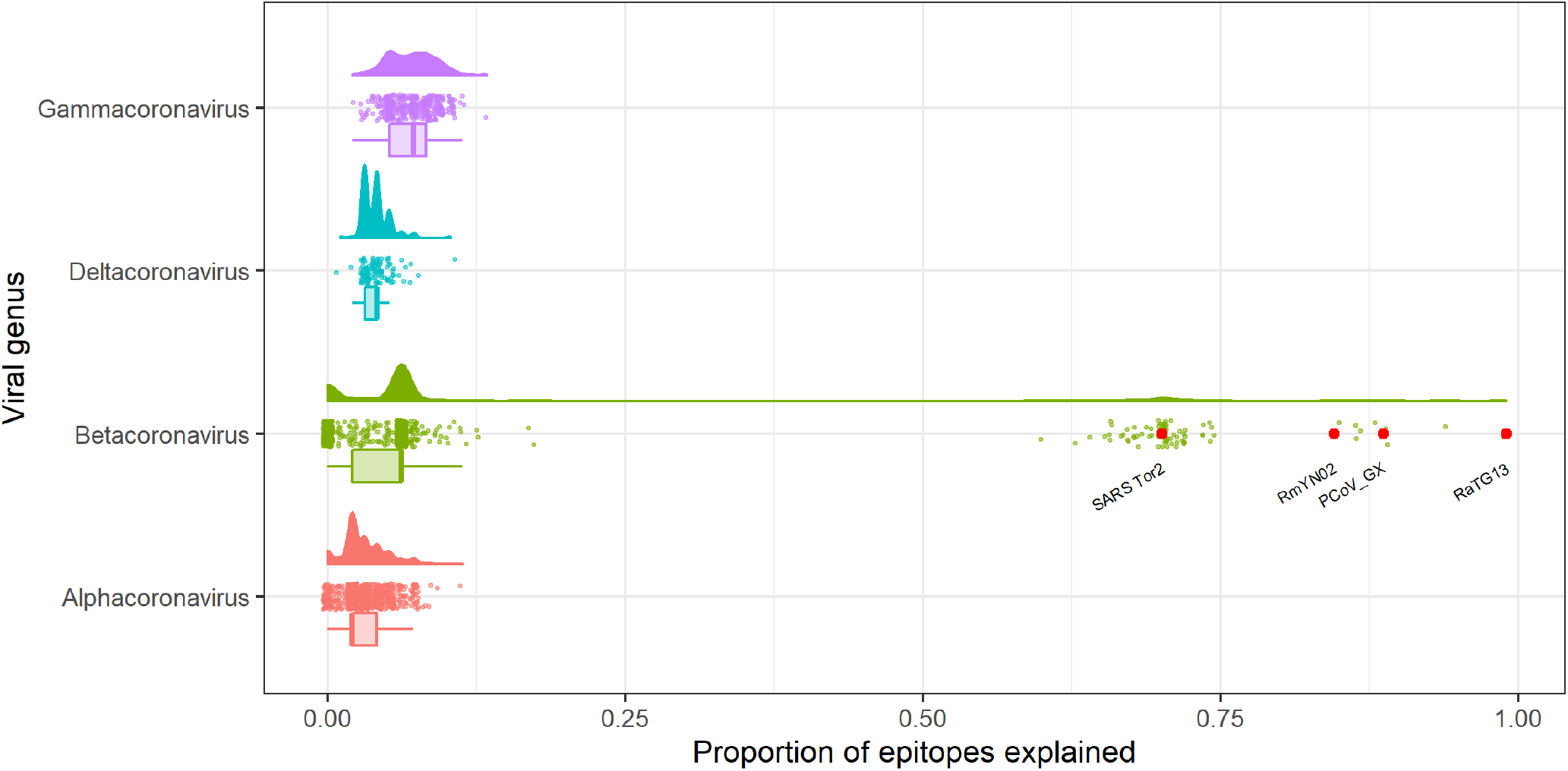
Proportion of ‘unexplained’ epitopes that have detectable sequence homology to members of Coronaviridae. Raincloud plot (57) of the proportion of ‘unexplained’ epitopes that have detectable homology to each coronavirus in our dataset (excluding SARS-CoV-2).

**Table S1. Curated metadata of the 2531 viral records in the *Coronaviridae***.

**Table S2. Proportion of epitopes with detectable homology to proteins of the *Coronaviridae***. (a) Proportion of 97 ‘unexplained’ epitopes explained by each of the viruses in our dataset (excluding HCoVs and SARS-CoV-2). (b) Proportion of all 177 published epitopes for 155 viruses with unique host and viral species (excluding SARS-CoV-2). These tables were generated using a custom *R* script (github.com/cednotsed/tcell_cross_reactivity_covid/blob/main/plot_deconvoluted_hcov_heatmap.R).

**Table S3. Protein BLAST results of 177 published epitopes against non-*Coronaviridae* proteins**. Merged protein BLAST output of eight searches (https://tinyurl.com/y22o4t9z). Merging was performed using a custom *R* script (github.com/cednotsed/tcell_cross_reactivity_covid/blob/main/utils/merge_web_blast.R).

**Table S4. GISAID acknowledgements table for the 12 bat and pangolin coronavirus sequences**.

**Table S5.** (a) List of 177 epitopes used in this study, including their respective study source and T-cell response type. (b) Frequency table generated from **Table S5a** stratified by study name and T-cell response type.

