## Supplementary material for "Pre-existing T cell-mediated cross-reactivity to SARS-CoV-2 cannot solely be explained by prior exposure to endemic human coronaviruses": Table S4

We gratefully acknowledge the following Authors from the Originating laboratories responsible for obtaining the specimens, as well as the Submitting laboratories where the genome data were generated and shared via GISAID, on which this research is based.

All Submitters of data may be contacted directly via www.gisaid.org

| Accession ID | Originating Laboratory | Submitting Laboratory | Authors |
|---|---|---|---|
| EPI_ISL_402131 | Wuhan Institute of Virology, Chinese Academy of Sciences | Wuhan Institute of Virology, Chinese Academy of Sciences | Yan Zhu, Ping Yu, Bei Li, Ben Hu, Hao-Rui Si, Xing-Lou Yang, Peng Zhou, Zheng-Li Shi |
| EPI_ISL_410538, EPI_ISL_410539, EPI_ISL_410540, EPI_ISL_410541, EPI_ISL_410542, EPI_ISL_410543, EPI_ISL_410544 | Beijing Institute of Microbiology and Epidemiology | Beijing Institute of Microbiology and Epidemiology | Wu-Chun Cao; Tommy Tsan-Yuk Lam; Na Jia; Ya-Wei Zhang; Jia-Fu Jiang; Bao-Gui Jiang |
| EPI_ISL_410721 | South China Agricultural University | South China Agricultural University | Yongyi Shen, Lihua Xiao, Wu Chen |
| EPI_ISL_412860 | SCSFRI, South China Sea Fisheries Research Institute, Chinese Academy of Fishery Sciences (SCSFRI, CAFS) | SCSFRI, South China Sea Fisheries Research Institute, Chinese Academy of Fishery Sciences (SCSFRI, CAFS) | Jiang,J.-Z., Liu,P. and Chen,J.-P. |
| EPI_ISL_412976, EPI_ISL_412977 | Shandong First Medical University & Shandong Academy of Medical Sciences | Institute of Microbiology, Chinese Academy of Sciences | Weifeng Shi, Tao Hu, Hong Zhou, Juan Li, Xing Chen, Alice Catherine Hughes, Yuhai Bi |
